# IBD-Derived Colonic Fibroblasts Exhibit an Osteopontin-Enriched Secretome, and Osteopontin Restrains Human Colonic Organoid Maturation

**DOI:** 10.64898/2026.08.28.747568

**Authors:** Dimitri Hamel, Muriel Quaranta, Delphine Bonnet, Laurent Alric, Audrey Ferrand

## Abstract

**Background:** Intestinal fibroblasts are extensively remodeled in inflammatory bowel disease (IBD), yet the soluble stromal signals that directly influence epithelial maturation remain incompletely understood. We examined whether fibroblasts derived from inflamed IBD colon display an osteopontin (OPN; SPP1)-enriched secretory phenotype and whether extracellular OPN directly modifies non-neoplastic human colonic epithelium.

**Methods:** Conditioned media from 5 noninflamed-associated fibroblast (NAF) and 4 inflammatory-associated fibroblast (IAF) cultures were analyzed in the validated multi-donor cytokine-array matrix, with orthogonal SPP1 RT-qPCR validation in a complementary fibroblast cohort. Recombinant OPN was then tested in human colonic organoids from 3 donors using donor-resolved molecular and functional analyses under standard, fibroblast-conditioned, and WNT-modified culture conditions. Donor identity defined biological replication.

**Results:** OPN showed the strongest positive rank-based separation between IAF and NAF cultures: all 4 IAF values were higher than all 5 NAF values (Cliff’s delta=1.00; exact Mann-Whitney P=.0159; median ratio=3.64; Benjamini-Hochberg q=.19). Fibroblast RT-qPCR showed approximately 10-fold higher mean SPP1 expression in IAF than NAF cultures (P<.05). In organoids, OPN consistently reduced KRT20, FABP1, CA2, and MUC2 from Day 5 to Day 9. SOX9, HES1, and NOTCH1 increased at Day 9, whereas LGR5 and ALDH provided no evidence of canonical stem-cell expansion. Organoid-area and EdU responses were modest and donor dependent.

**Conclusions:** IBD-derived colonic fibroblasts can display an OPN-enriched secretory phenotype. In human colonic organoids, OPN is sufficient to impair epithelial maturation, whereas its effects on growth and proliferation are variable and depend on the surrounding niche.

**Key Messages:** *What is already known?:* IBD remodels intestinal fibroblasts, however the direct effects of fibroblast-associated osteopontin on human colonic epithelial maturation are poorly defined.

*What is new here?:* In a validated multi-donor NAF/IAF matrix, OPN was the only analyte with complete positive separation between groups, was supported by SPP1 RT-qPCR, and induced sustained loss of maturation markers followed by a later SOX9/HES1/NOTCH1-associated state in human colonoids.

*How can this study help patient care?:* The data identify stromal OPN as a candidate contributor to incomplete epithelial normalization in IBD and provide a human experimental framework for testing whether endogenous OPN blockade can restore epithelial maturation.

## Introduction

Inflammatory bowel diseases arise from sustained interactions between the immune system, the stromal compartment, and the intestinal epithelium. Single-cell studies of the human intestine have highlighted how chronic inflammation remodels the colonic mesenchyme, with the emergence or expansion of activated fibroblast populations characterized by altered extracellular matrix production and paracrine signaling (1-4). Fibroblasts are therefore no longer considered simply as bystanders to inflammation, but as active contributors to the persistence and organization of the diseased mucosal microenvironment.

Epithelial repair is similarly dynamic. Following injury, intestinal epithelial cells transiently acquire more plastic states before progressively re-establishing mature absorptive and secretory functions. However, both human studies and experimental models suggest that these injury-associated or inflammatory epithelial states can persist when the surrounding niche remains altered (5-7). Identifying the stromal signals that interfere with this transition may therefore help explain why epithelial architecture and function are not always fully restored even when overt inflammation is controlled.

Osteopontin (OPN), encoded by *SPP1*, is a secreted phosphoglycoprotein involved in cell adhesion, immune responses, and tissue remodeling. Increased circulating OPN concentrations are observed in patients with IBD and have been associated with disease activity (8,9). However, OPN functions in the intestine appear to be highly dependent on the context. OPN has been implicated in inflammatory-cell recruitment and innate defense, but also in epithelial hyperplasia and tissue repair (9-11). More recently, macrophage-derived SPP1, together with NRG1, was identified as part of a regenerative niche that promotes injury-associated epithelial reprogramming in mouse and human organoid models (12). OPN produced by myofibroblasts has also been implicated in colitis-associated tumorigenesis, further suggesting that stromal cells may represent an important local source of OPN in the diseased intestine (13).

In consequence, we investigated whether primary fibroblasts derived from inflamed human IBD colon display an OPN-enriched secretory phenotype and, independently, whether OPN can directly influence the state of non-neoplastic human colonic epithelium. We first examined OPN within a validated multi-donor cytokine-array dataset and then assessed the epithelial response to recombinant OPN in normal human colonic organoids using donor-resolved analyses. Rather than considering organoid growth as a single endpoint, we separately examined maturation, stem-cell-associated programs, proliferation, and growth. Finally, we tested OPN under fibroblast-conditioned and WNT-modified culture conditions to determine whether its epithelial effects are shaped by the surrounding niche.

## Materials and Methods

### Human Tissue, Primary Fibroblasts, and Colonic Organoids

Human colonic tissues were obtained at Toulouse University Hospital through the registered COLIC collection (ethics approval DC-2015-2443) approved by the French South-West and Overseas IV ethics committee. Patients provided informed consent before sample collection. Primary fibroblast cultures were established from macroscopically noninflamed colonic tissue (NAF) or inflamed IBD tissue (IAF). The IAF bank comprised 3 Crohn disease cultures (IAF1, IAF2, IAF3) and 1 ulcerative colitis culture (IAF4). The validated cytokine-array matrix contained 5 NAF cultures (NAF1, NAF2, NAF3, NAF4, NAF5) and all 4 IAF cultures. NAF4 was also used to generate conditioned medium for complementary niche experiments. Normal human colonic organoid cultures used for the principal donor-aware analyses were NORG1, NORG2, and NORG3. Clinical information is reported in Supplementary Table S1.

### Fibroblast Establishment and Culture Quality Control

Fibroblasts were isolated from colonic stromal tissue after epithelial crypt removal and collagenase IV digestion and expanded in minimum essential medium (MEM) supplemented with fetal bovine serum. Mesenchymal identity and culture purity were assessed during primary-culture establishment. The final inferential fibroblast analyses reported here are restricted to the retained NAF/IAF secretome cohort and the complementary SPP1 RT-qPCR validation cohort.

### Fibroblast Conditioned Medium and Cytokine Array

Fibroblasts were cultured to approximately 70%-80% confluence, switched to minimum essential medium containing 0.5% fetal bovine serum, and conditioned medium was collected after 48 hours, centrifuged, and stored at -80°C when used for secretome analysis. Soluble factors were measured with the Human XL Cytokine Array (R&D Systems, ARY022B) according to the manufacturer’s protocol. Chemiluminescent spot intensities were quantified with ImageJ. Secretome analysis used the validated publication matrix comprising 5 biologically independent NAF cultures and 4 IAF cultures processed within the same harmonized normalization workflow. Thirty-six analytes had complete normalized values across all 9 cultures. Analytes were ranked with Cliff’s delta; nominal comparisons used exact two-sided Mann-Whitney tests, with Benjamini-Hochberg correction across the complete 36-analyte panel. Leave-one-donor-out analyses were performed for OPN. A Crohn-only sensitivity analysis used IAF1, IAF2, and IAF3.

### Organoid Culture and OPN Exposure

Colonic crypts were embedded in Matrigel and expanded using a WENR-based human colonic organoid medium adapted from Sato et al. (14). Recombinant human OPN (PeproTech, 120-35) was tested at 0.1, 1, and 10 μg/mL in dose-response experiments. Principal WENR experiments used 10 μg/mL OPN with matched vehicle. The 10 μg/mL concentration was selected empirically from the experimental dose-response and is not presented as a physiological tissue concentration.

### Niche-Perturbation Experiments

To test niche dependence, organoids were also cultured in NAF4-conditioned medium and, in independent experiments, in media containing either 25% or 1% (v/v) WNT3A-conditioned medium. The percentages refer to the volumetric fraction of laboratory-produced WNT3A-conditioned medium, not to a molecular WNT3A concentration. Within each experiment, OPN-treated cultures were compared with the corresponding vehicle control. Independent culture contexts were not treated as paired conditions.

### High-Content Organoid Analysis, EdU, and ALDH

Bright-field images were acquired using the Opera Phenix platform and analyzed with an automated pipeline to identify organoids and quantify projected area and morphology. Organoid-level observations were treated as nested within-donor measurements and not as independent biological replicates. Day 9 proliferation was assessed by 1-hour incorporation of 15 μmol/L 5-ethynyl-2’-deoxyuridine (EdU), followed by Click-iT detection, DAPI counterstaining, and confocal high-content imaging. ALDH activity was measured with the ALDEFLUOR assay using diethylaminobenzaldehyde as the negative control.

### RNA Extraction and RT-qPCR

RNA was extracted with Direct-zol (Zymo Research), and 500 ng RNA was reverse transcribed using the Maxima First Strand cDNA Synthesis Kit. Quantitative PCR was performed on a LightCycler 480 using Takyon SYBR MasterMix, 600 nM gene-specific primers, and 5 ng cDNA per reaction. Reactions were run in technical triplicate. All primer assays used in the study had validated efficiencies between 90% and 110%, and specificity was verified by melt-curve analysis. Expression was normalized to GAPDH, B2M, and HPRT after verifying stability of the reference genes under vehicle and OPN conditions. Primer sequences are provided in Supplementary Table S3 and reporting follows MIQE principles (17). Technical triplicates were averaged before biological analysis. Statistical testing was performed on DeltaCq-derived normalized quantities, whereas fold changes relative to matched vehicle are used for graphical representation. Fibroblast SPP1 validation was performed in the 5 NAF and 4 IAF cultures and compared with a two-sided unpaired Student t test on normalized qPCR values.

### Statistical Analysis

The patient-derived culture was the biological experimental unit. Individual organoids, nuclei, and technical qPCR wells were not counted as independent biological replicates. For WENR qPCR, donor-resolved trajectories were summarized using a predefined differentiation-associated module (KRT20, FABP1, CA2, MUC2) and an immature/NOTCH-associated module (SOX9, HES1, NOTCH1), calculated as geometric means of fold changes for visualization. In NAF4-conditioned medium, the same differentiation module was retained; complementary summaries used an immaturity/NOTCH module (SOX9, HES1, NOTCH1, CD133) and a KI67/CD44 module (KI67, CD44v1). These module summaries were descriptive at n=3 and were not used for significance fishing. Day 9 organoid-area effects were summarized as the OPN/vehicle median-area ratio per donor. Day 9 EdU was summarized as the percentage of EdU-positive nuclei per donor. WNT3A-conditioned-medium interaction analyses used donor-level OPN/vehicle response ratios; the 1% versus 25% interaction ratio was 0.966 for area (95% CI, 0.790-1.181; P=.534) and 0.793 for EdU (95% CI, 0.401-1.567; P=.280). Given the small biological n, exact values, effect sizes, directionality, and uncertainty were prioritized over significance-star reporting. Statistical analysis were performed using GraphPad Prism 11, and with support from OpenAI ChatGTP.

### Contextualization With Published Human Crohn Disease Data

Published human datasets were used to interpret the experimental findings, not as a substitute for de novo validation. We prioritized Crohn disease studies because 3 of the 4 IAF cultures were Crohn derived. No de novo pseudobulk reanalysis of public single-cell matrices is claimed.

## Results

### A Validated Multi-Donor Secretome Matrix Identifies OPN as the Strongest Positive IAF-Associated Signal

Across the nine fibroblast cultures, 36 analytes had complete normalized values. As expected from primary human samples, the overall secretome profiles were heterogeneous. Despite this variability, OPN emerged as the most consistent IAF-associated signal. Indeed, it was the only analyte for which all 4 IAF values were higher than all 5 NAF values, resulting in complete positive rank separation between the two groups (Cliff’s delta=1.00; Figure 1A-B). The median OPN signal was 3.64-fold higher in IAF than in NAF cultures, with an exact two-sided Mann-Whitney P value of 0.0159 (Figure 1C). This association did not remain significant after correction for multiple testing across the 36-analyte panel (Benjamini-Hochberg q=0.19).

**Figure 1.**
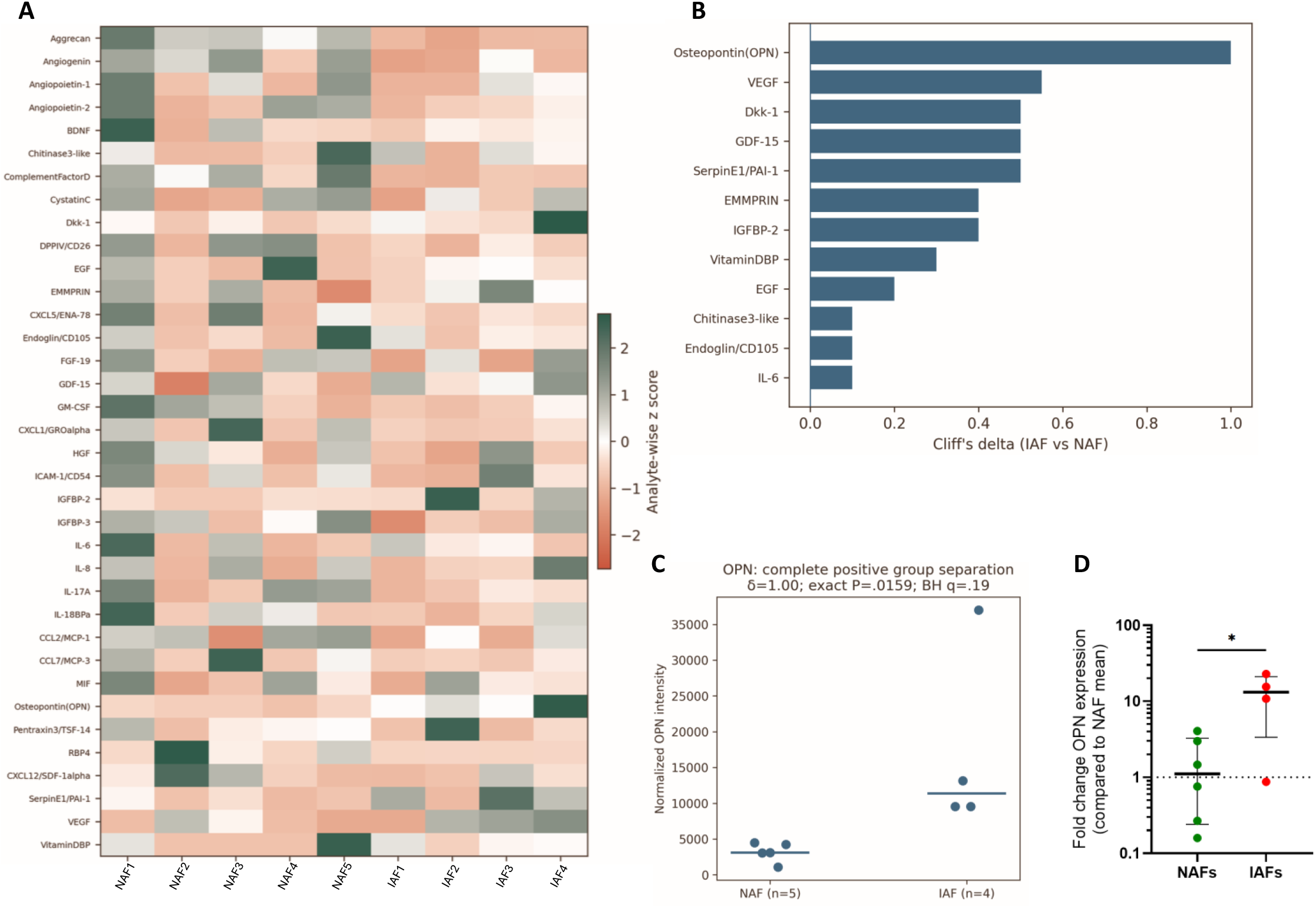
A validated multi-donor cytokine-array matrix identifies OPN as the strongest positive IAF-associated secretome signal and is supported by orthogonal SPP1 RT-qPCR. (A) Analyte-wise z-score heatmap of 36 factors across 5 NAF and 4 IAF cultures. The heatmap is centered at zero: negative z scores progress from white to red and positive z scores from white to dark green. (B) Leading positive IAF-associated analytes ranked by Cliff’s delta. OPN is the only analyte with complete positive group separation. (C) Donorlevel normalized OPN intensities. Horizontal bars indicate group medians; all 4 IAF values exceed all 5 NAF values (Cliff’s delta=1.00; exact two-sided Mann-Whitney P=.0159; median IAF/NAF ratio=3.64; Benjamini-Hochberg q=.19). (D) Orthogonal SPP1 RT-qPCR validation in a complementary 5-NAF/4-IAF fibroblast cohort; mean expression was approximately 10-fold higher in IAF cultures (two-sided unpaired Student t test, P<.05).

We therefore interpreted OPN as the leading candidate emerging from this exploratory screen, rather than as a statistically confirmed discovery after FDR correction.

### The OPN Signal Is Robust to Donor Removal and Is Supported at the Transcript Level

The complete separation between IAF and NAF cultures was not driven by a single donor, as it was preserved in all leave-one-donor-out analyses. This pattern also remained evident when the analysis was restricted to the 3 Crohn disease-derived IAF cultures (IAF1, IAF2, IAF3). All of them showed higher OPN levels than the 5 NAF cultures (Cliff’s delta=1.00; exact P=0.0357; median IAF/NAF ratio=4.22; Supplementary Figure S1).

We then assessed the expression of SPP1 mRNA, coding for OPN, independently by RT-qPCR in the same 5 NAF and 4 IAF cultures. Mean SPP1 expression was approximately 10-fold higher in IAF than in NAF cultures (P<0.05; Figure 1D), providing transcript-level support for the secretome finding. Together, these results consistently point to an OPN/SPP1-enriched phenotype in IAF cultures, while the exploratory nature and limited size of the secretome dataset should still be kept in mind.

### OPN Induces an Early and Persistent Suppression of Epithelial Maturation Programs

Normal colonic organoids were cultured in WENR medium in the presence of 10 μg/mL recombinant OPN. Across donors, the clearest and most consistent effect of OPN was observed at the transcriptional level. Expression of the differentiation markers KRT20, FABP1, CA2, and MUC2 was already reduced in most donors at Day 5 and became consistently lower across NORG1, NORG2, and NORG3 by Day 9 (Figure 2A). At Day 9, the geometric-mean fold changes were approximately 0.62 for KRT20, 0.28 for FABP1, 0.48 for CA2, and 0.26 for MUC2. Consistent with these individual markers, the predefined differentiation-associated module remained clearly below vehicle levels at both time points (Figure 2B), indicating that OPN produces an early and sustained impairment of epithelial maturation.

**Figure 2.**
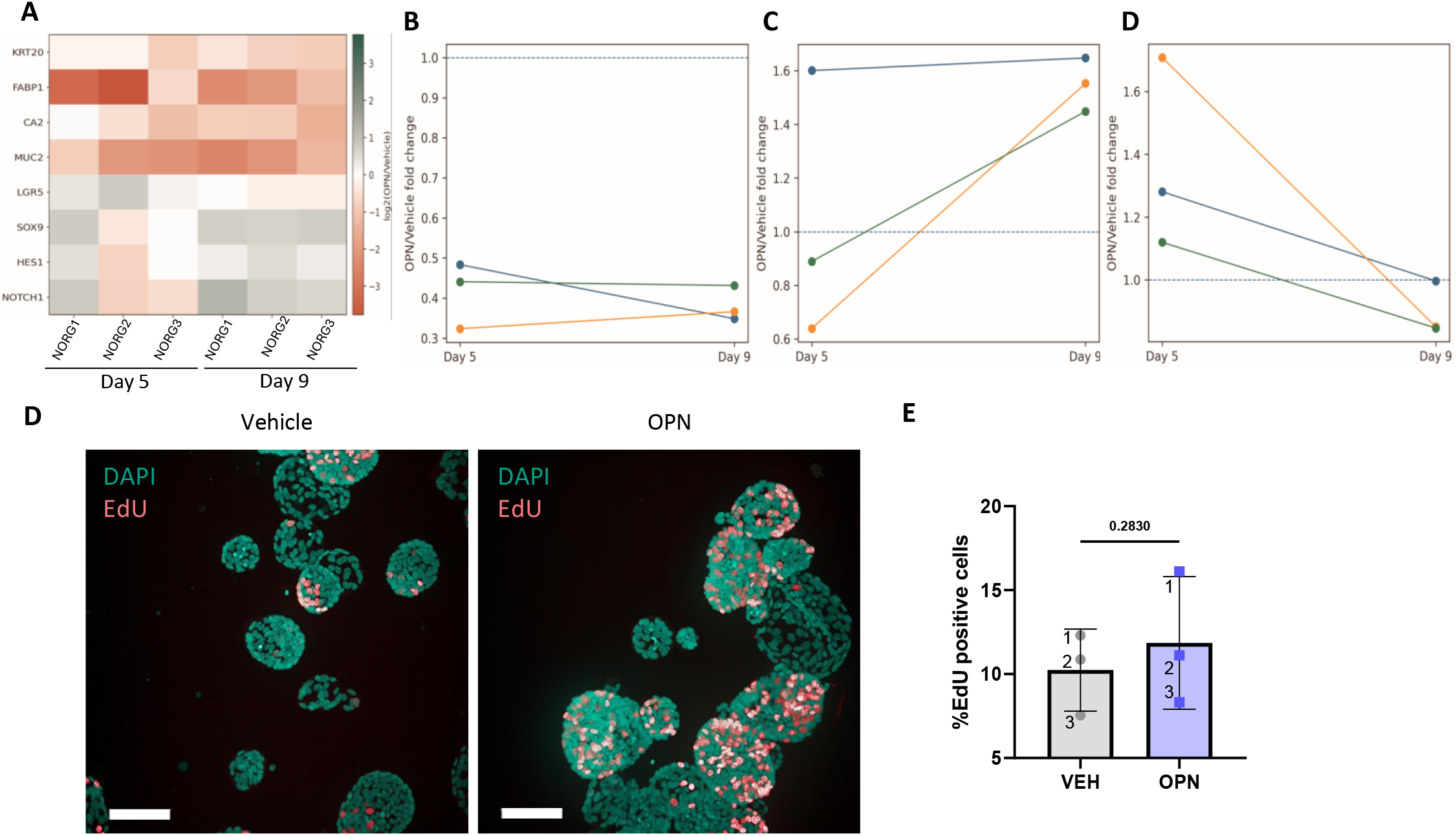
OPN produces an early loss of maturation-associated transcription followed by a later immature/NOTCH-associated state. (A) Donor-resolved log2 fold changes for KRT20, FABP1, CA2, MUC2, LGR5, SOX9, HES1, and NOTCH1 at Day 5 and Day 9 in WENR after OPN relative to matched vehicle. The heatmap is centered at zero with negative values in red and positive values in green. (B) Differentiation-associated module. (C) Immature/NOTCH-associated module. (D) LGR5. Lines in B-D connect Day 5 and Day 9 values from the same donor. (E) Day 9 donor-level OPN/vehicle median-area ratios; donors are independent and are not connected. (F) Day 9 EdU-positive nuclei; lines connect matched vehicle and OPN values within each donor.

### A Later Immature/NOTCH-Associated Program Emerges Without Sustained LGR5 Expansion

SOX9, HES1, and NOTCH1 showed more variable responses at Day 5, but by Day 9 all three were increased across the 3 principal organoid donors. Their geometric-mean fold changes at Day 9 were approximately 1.64, 1.27, and 1.78, respectively (Figure 2A, C). In contrast, LGR5 was not maintained above the levels observed for vehicle condition at Day 9 (geometric mean approximately 0.89; Figure 2D). Moreover, we did not observe a reproducible increase in the proportion of ALDH-high epithelial cells (Data not shown).

These results argue against a simple expansion of the canonical LGR5-positive stem-cell compartment. Instead, they suggest that OPN first interferes with epithelial differentiation and is then associated with the emergence of a more immature, NOTCH-associated transcriptional state.

### Growth and Proliferation Responses Are Modest and Donor Dependent

At Day 9, the effect of OPN on organoid growth was modest and varied between donors. The OPN/vehicle median-area ratios were 0.94 for NORG1, 1.18 for NORG2, and 1.19 for NORG3, corresponding to an overall geometric mean of approximately 1.10 (exploratory P=.343; Figure 2E). A similar pattern was observed for proliferation. The proportion of EdU-positive nuclei increased from 12.3% to 16.1% in NORG1, from 10.9% to 11.1% in NORG2, and from 7.5% to 8.3% in NORG3, but this change was not significant at the donor level (paired P=0.282; Figure 2F). Because individual organoids and nuclei are nested within each donor, we based the statistical interpretation on donor-level summaries rather than on pooled object-level measurements.

Overall, these results suggest that the effect of OPN on proliferation is limited and donor dependent, whereas the impairment of epithelial maturation is the most consistent response observed across the organoid cultures.

### The Epithelial Response to OPN Is Niche Dependent

The epithelial response to OPN treatment also depended on the surrounding culture environment. Based on the observation of the secretome matrix, we treated the epithelial cultures with NAF4-conditioned medium. In this condition, the differentiation-associated module showed no consistent direction of change across donors at Day 15. In contrast, the immaturity/NOTCH-and KI67/CD44-associated summaries increased in all 3 organoid lines (Supplementary Figure S3). This was particularly evident for KI67, which increased 1.45-, 2.63-, and 1.31-fold, and for CD44v1, which increased 1.24-, 2.20-, and 1.34-fold in NORG1, NORG2, and NORG3, respectively. The broader Fluidigm dataset pointed in the same direction, with increased expression of genes related to cell cycle, NOTCH signaling, epithelial plasticity, and candidate OPN receptors, while canonical intestinal stem-cell markers remained comparatively stable. However, because a complete donor-resolved matrix was not available for every target in this extended panel due to sample issues, these results were considered descriptive.

We then asked whether modifying WNT availability would further influence the epithelial response to OPN. Under the 1% WNT3A-conditioned-medium condition, KRT20 and FABP1 decreased in all 3 donors, whereas NOTCH1 showed a modest increase in each donor. Responses were less consistent in the 25% WNT3A-conditioned-medium condition (Figure 3). Despite these transcriptional differences, reducing the proportion of WNT3A-conditioned medium did not measurably enhance the effects of OPN on organoid growth or proliferation. The 1%/25% interaction ratio was 0.966 for organoid area (95% CI, 0.790-1.181; P=.534) and 0.793 for EdU incorporation (95% CI, 0.401-1.567; P=.280; Supplementary Figure S4).

**Figure 3.**
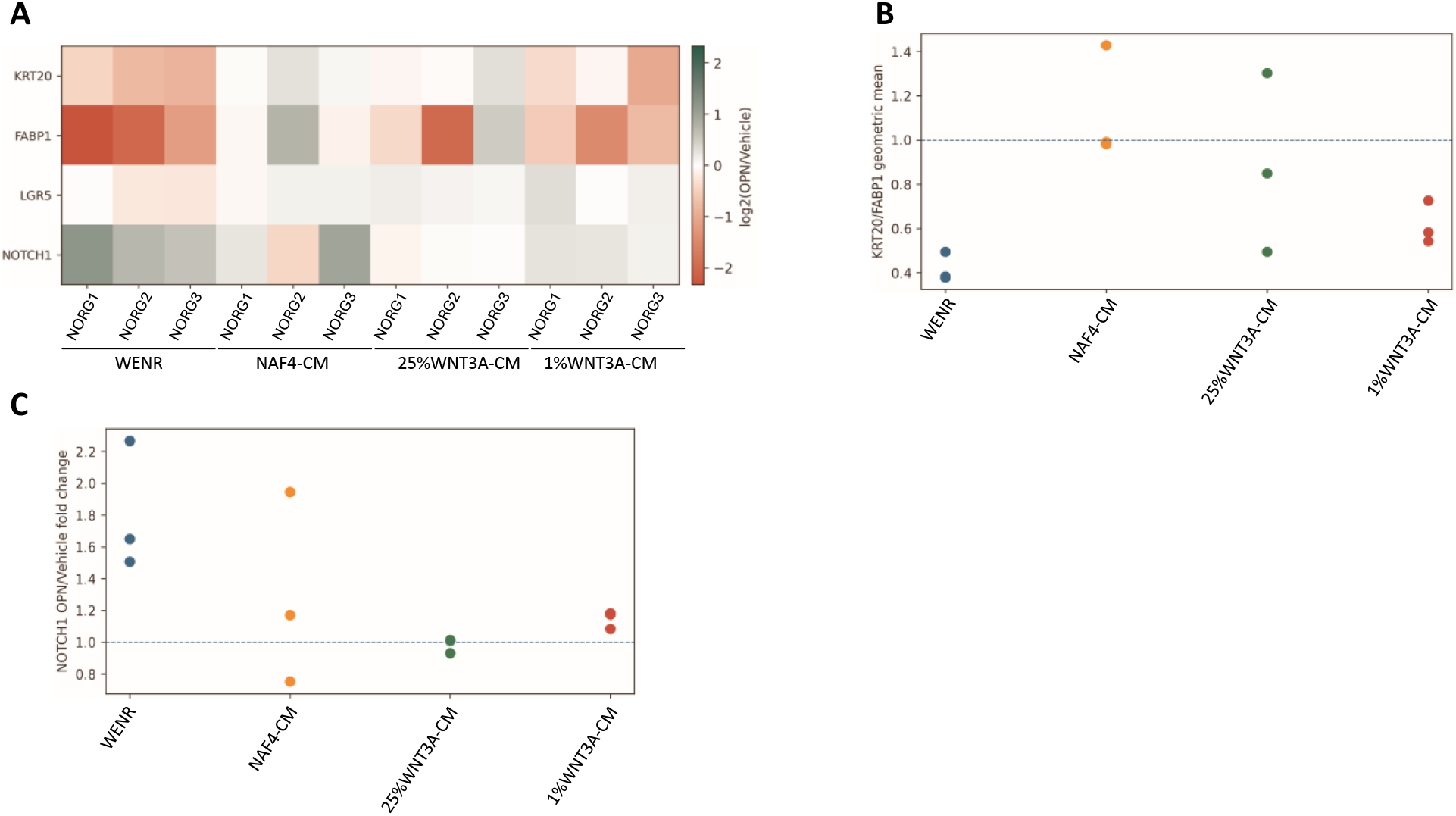
The epithelial transcriptional response to OPN depends on the surrounding niche. (A) Donorresolved log2 fold changes for KRT20, FABP1, LGR5, and NOTCH1 in WENR, NAF4-conditioned medium, 25% WNT3A-conditioned medium, and 1% WNT3A-conditioned medium. (B) KRT20/FABP1 differentiation-module values. (C) NOTCH1 values. Independent culture contexts are shown as unconnected donor-level points and are not treated as a single matched factorial experiment.

Together, these results indicate that the transcriptional response to OPN is shaped by the surrounding niche, although in our experimental conditions, this context dependence did not translate into a clear functional interaction for growth or proliferation.

## Discussion

Our results point to OPN as a prominent soluble feature of inflammatory IBD-derived colonic fibroblasts and, importantly, show that extracellular OPN can directly alter the maturation state of non-neoplastic human colonic epithelium. The epithelial response was much more consistent at the transcriptional level than at the level of growth or proliferation. Differentiation-associated genes were reduced early and remained suppressed, whereas SOX9, HES1, and NOTCH1 increased at later time points. In contrast, LGR5 was not maintained above control levels and the ALDH-high epithelial fraction did not expand. Taken together, these findings suggest that OPN primarily modifies epithelial cell state rather than acting as a general mitogenic factor.

This fibroblast phenotype fits with the broader remodeling of the IBD stroma described in human tissue. Kinchen et al. showed that colitis disrupts homeostatic crypt-niche mesenchymal populations and promotes the emergence of activated stromal states with altered epithelial-support functions (1). Subsequent studies have further highlighted the extent of fibroblast heterogeneity and disease-associated stromal communication in Crohn disease (2,3). Our data do not demonstrate that OPN is the dominant stromal mediator in vivo. They do, however, identify OPN as a soluble factor produced at higher levels by inflammatory primary human fibroblasts and show that it is capable of directly modifying the phenotype of human colonic epithelium.

The role of OPN in intestinal inflammation and repair is likely to be more complex than a simple deleterious or protective effect. OPN has been associated with inflammatory-cell recruitment and disease activity in IBD (8-10), but it has also been implicated in epithelial responses to infection and tissue injury (11,12). The study by Moraitis et al. is particularly relevant in this respect, because macrophage-derived SPP1 was identified together with

NRG1 as part of a regenerative niche rather than as an isolated instructive signal (12). Our findings are consistent with such a context-dependent role. Recombinant OPN alone was sufficient to delay epithelial maturation, but the phenotype induced by OPN changed substantially when the composition of the surrounding fibroblast- or WNT-derived niche was modified.

The experiments performed in NAF-conditioned medium illustrate this point. Under these conditions, the strong and consistent differentiation defect observed in WENR was no longer reproduced across all donors. Instead, the KI67/CD44- and immaturity/NOTCH-associated signatures tended to increase. The broader Fluidigm panel supported the same general trend, although these data should remain descriptive because a complete donor-resolved numerical matrix was not available for every target. Likewise, reducing the proportion of WNT3A-conditioned medium modified the transcriptional response to OPN, but did not result in a statistically supported OPN-by-WNT interaction for either organoid area or EdU incorporation. These observations therefore argue against a single, context-independent mechanism through which OPN would uniformly stimulate epithelial proliferation.

Several limitations should be considered when interpreting these findings. First, the secretome analysis is based on a single harmonized multi-donor cytokine-array matrix comprising 5 NAF and 4 IAF cultures. OPN completely separated the two groups and remained the strongest positive IAF-associated signal in leave-one-donor-out analyses, but the comparison did not remain significant after correction for multiple testing. Complete separation was also preserved when the analysis was restricted to the 3 Crohn disease-derived IAF cultures, although this small subgroup is clearly insufficient to establish a Crohn-specific secretory phenotype. Second, the epithelial experiments were performed using 3 principal organoid donors, and the molecular phenotype was assessed mainly by RT-qPCR. Most importantly, the use of recombinant OPN demonstrates sufficiency, but not necessity. We have not yet determined whether neutralizing OPN, or reducing endogenous SPP1 in IAF-conditioned medium, is sufficient to reverse the epithelial phenotype. This will be essential to establish a causal fibroblast-OPN-epithelium axis. Finally, the NAF cultures were obtained from macroscopically noninflamed colonic tissue rather than from uniformly healthy donors, and several were derived from resections performed for non-IBD diseases.

Despite these limitations, the combination of a donor-defined fibroblast secretome signal with a reproducible effect on epithelial maturation supports a model in which inflammatory stromal remodeling may increase epithelial exposure to OPN and thereby contribute to the persistence of an incompletely mature epithelial state. Demonstrating the dependence of this phenotype on endogenous fibroblast-derived OPN, and defining the epithelial populations that respond to OPN at single-cell resolution, will be important next steps to establish the mechanistic and potential therapeutic relevance of this pathway.

## Conclusion

Primary fibroblasts isolated from inflamed IBD colon showed an OPN-enriched secretory profile in our multidonor cytokine-array screen. In normal human colonic organoids, OPN consistently reduced the expression of maturation-associated genes and was followed by the emergence of a more immature, NOTCH-associated transcriptional state. In contrast, its effects on organoid growth and proliferation were modest and varied between donors. Together, these findings identify OPN as a candidate stromal regulator of epithelial state in IBD and suggest that its effects are strongly influenced by the surrounding tissue niche.

## Data Availability

Data can be provided upon reasonable request to the corresponding author. Public datasets used for contextualization are GSE90607, GSE164985, and GSE266546 (18-20); no de novo pseudobulk analysis of these public matrices is claimed.

## Funding

Plan Cancer “System biology” 2017 C20048BS and INSERM. The region Occitanie and Université de Toulouse III funded the salary of D.H.

## Conflicts of Interest

The author declared no conflict of interest.

## Author Contributions

D. Hamel established the organoids and fibroblasts cultures, performed the experiment, analyzed the data, and prepared the manuscript.

M. Quaranta established the organoids and fibroblasts cultures

L. Alric and D. Bonnet screened the patients; received their consent, performed the endoscopy and the tissue biopsies.

A. Ferrand conceived and supervised the project, obtained the fundings, analyzed the data and wrote the manuscript.

## Acknowledgments

The authors thank the patients who consent in giving their tissues for our research. The authors thank staff from the INFINITy imaging platform for assistance with confocal imaging, as well as from the IRSD platform for assistance with HCS. Use of AI-assisted tools: OpenAI ChatGPT was used during manuscript preparation for language refinement, assistance with data-processing code, and figure-layout preparation. All numerical results, analyses, interpretations, references, and final text were critically reviewed and verified by the authors, who take full responsibility for the content of the manuscript.

## Use of AI-Assisted Tools

OpenAI ChatGPT was used during manuscript preparation for language refinement, assistance with statistical/data-processing code, and figure-layout preparation. All numerical results, analyses, interpretations, references, and final text were critically reviewed and verified by the authors, who take full responsibility for the content of the manuscript.

## Ethical Considerations

Human tissue collection was performed at Toulouse University Hospital through the registered COLIC collection (ethics approval DC-2015-2443) approved by the French South-West ethics committee. Patients provided informed consent before sample collection. Samples and study data were handled in deidentified form for the analyses reported here.

---

**Table 1.**
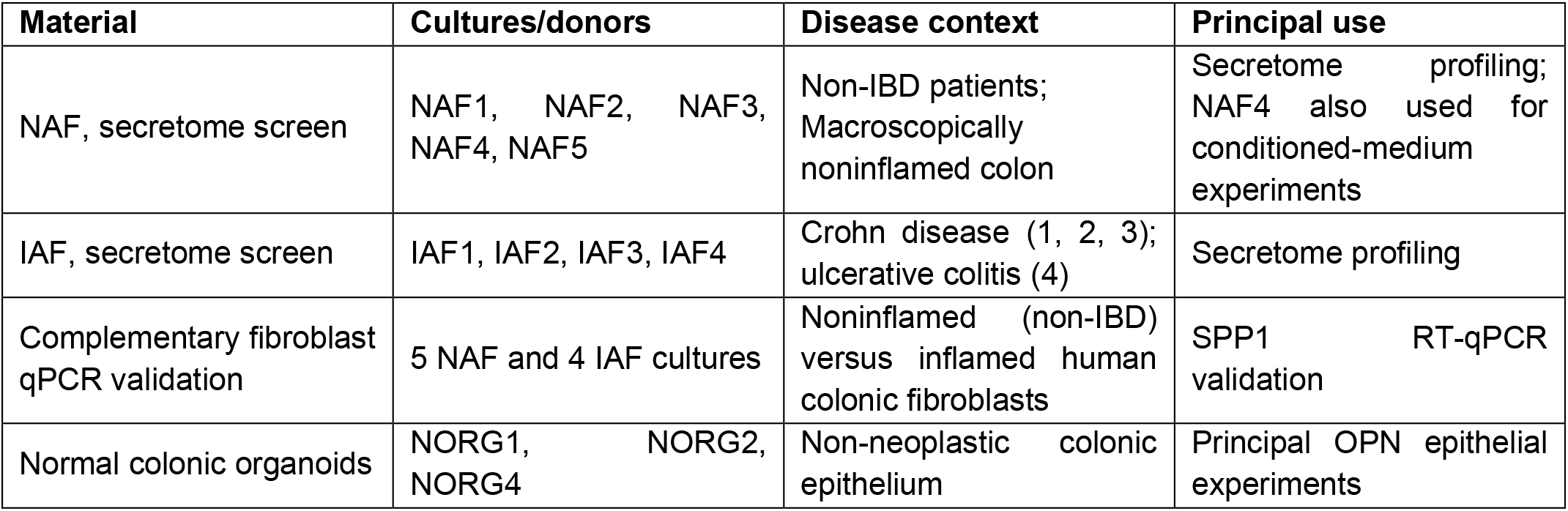
Biological Materials Used in the Principal Analyses.

| **Material** | **Cultures/donors** | **Disease context** | **Principal use** |
| --- | --- | --- | --- |
| NAF, secretome screen | NAF1, NAF2, NAF3, NAF4, NAF5 | Non-IBD patients; Macroscopically noninflamed colon | Secretome profiling; NAF4 also used for conditioned-medium experiments |
| IAF, secretome screen | IAF1, IAF2, IAF3, IAF4 | Crohn disease (1, 2, 3); ulcerative colitis (4) | Secretome profiling |
| Complementary fibroblast qPCR validation | 5 NAF and 4 IAF cultures | Noninflamed (non-IBD) versus inflamed human colonic fibroblasts | SPP1 RT-qPCR validation |
| Normal colonic organoids | NORG1, NORG2, NORG4 | Non-neoplastic colonic epithelium | Principal OPN epithelial experiments |

## Notes

### Competing Interest Statement

The authors have declared no competing interest.

