## Supplementary material for "IBD-Derived Colonic Fibroblasts Exhibit an Osteopontin-Enriched Secretome, and Osteopontin Restrains Human Colonic Organoid Maturation"

#### Supplementary Figure S1. Sensitivity analyses for the validated multi-donor secretome screen

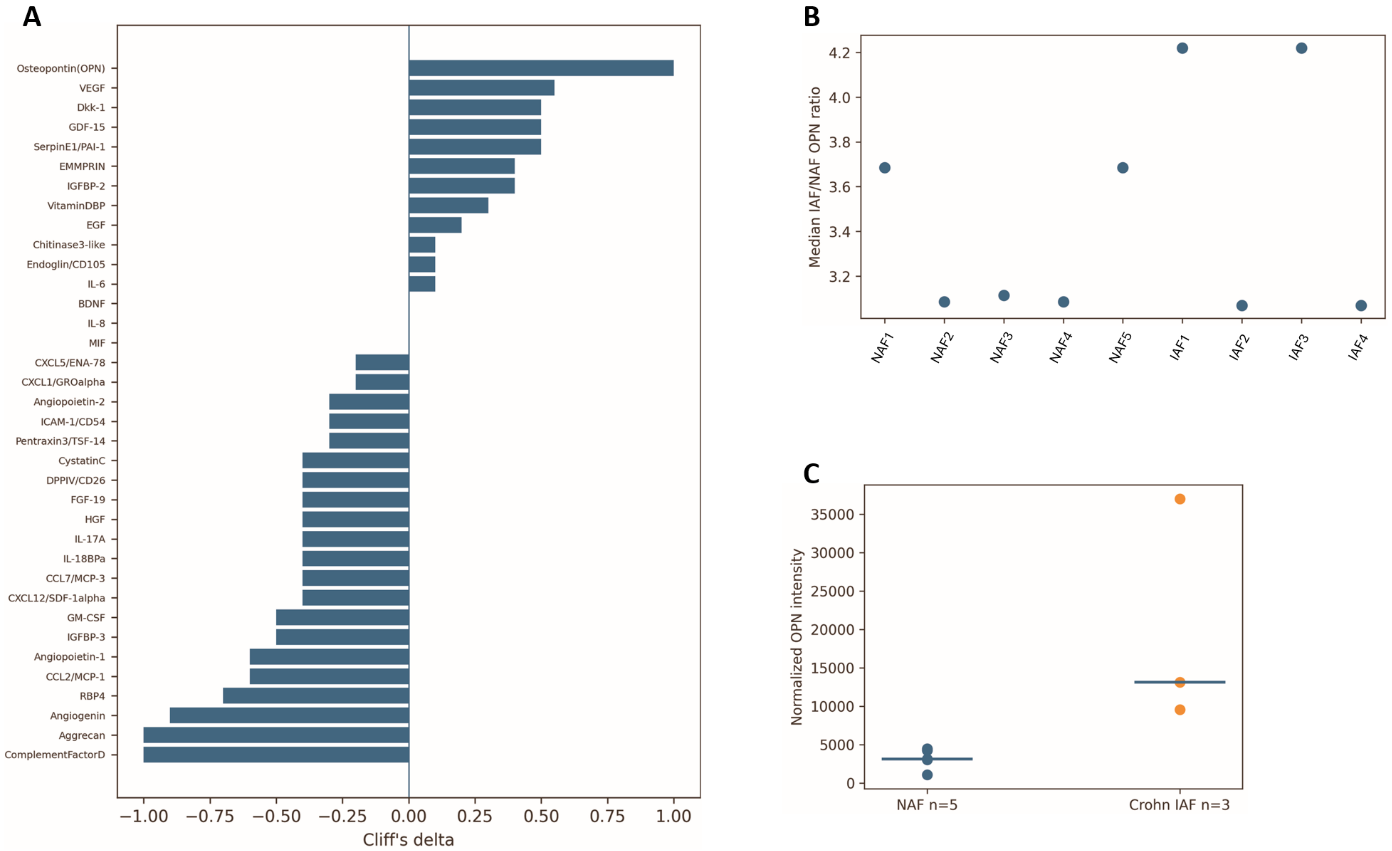


**Supplementary Figure S1.**

(A) Complete Cliff's-delta ranking for the 36 analytes with complete normalized values across 5 NAF and 4 IAF cultures. OPN is the only analyte showing complete positive group separation (delta=1.00).

(B) Leave-one-donor-out analysis of OPN; complete IAF-versus-NAF rank separation is preserved after removal of any single retained donor.

(C) Crohn-only sensitivity analysis using IAF1, IAF2, and IAF3 versus the 5 NAF cultures; complete separation is retained (delta=1.00; exact P=.0357; median ratio=4.22).

### Supplementary Figure S2. Complementary organoid functional analyses

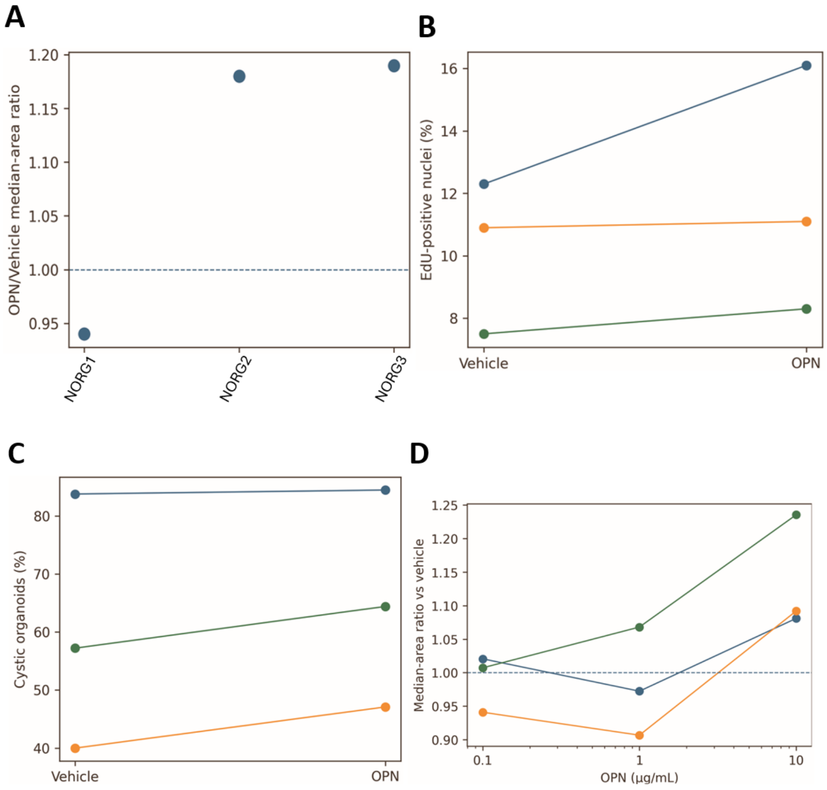


**Supplementary Figure S2.**

(A) Day 9 donor-level OPN/vehicle area ratios. Each point is an independent donor and points are not connected.

(B) Day 9 EdU-positive nuclei; lines connect matched vehicle and OPN values within each donor.

(C) Day 9 cystic morphology; lines connect paired vehicle and OPN values within each donor.

(D) Day 15 recombinant OPN dose response in NAF4-conditioned medium. Organoid-level observations are nested within donors and are not treated as biological n.

### Supplementary Figure S3. NAF4-conditioned-medium RT-qPCR and expanded Fluidigm response

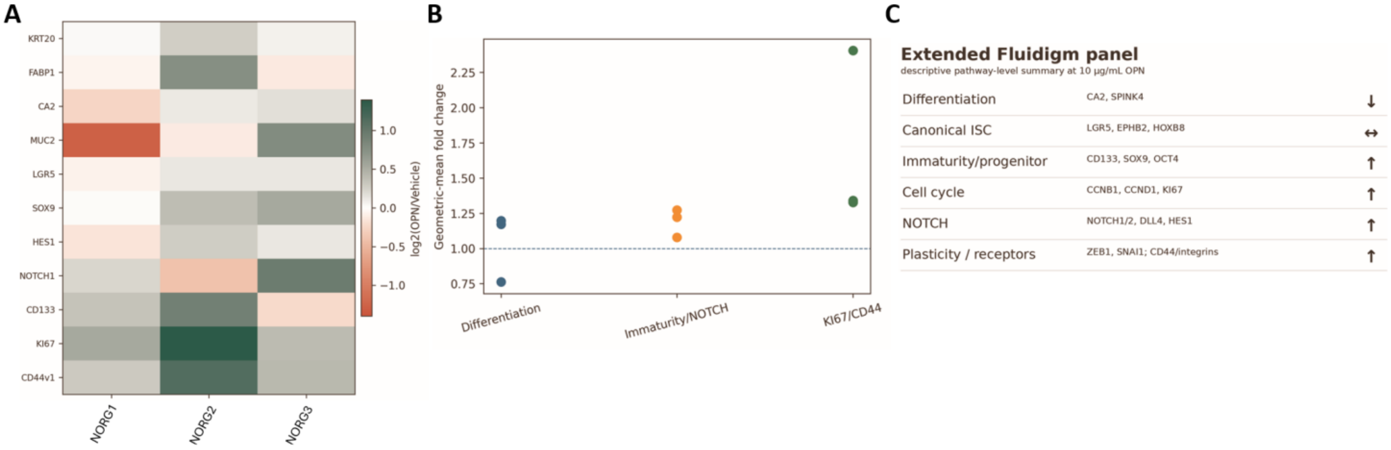


**Supplementary Figure S3.**

(A) Donor-resolved log2 OPN/vehicle fold changes at Day 15 for the quantitatively audited 11-gene NAF4-conditioned-medium panel. The heatmap is centered at zero, with negative values from white to red and positive values from white to dark green.

(B) Donor-level geometric-mean module summaries. The differentiation module comprises KRT20, FABP1, CA2, and MUC2; the immaturity/NOTCH module comprises SOX9, HES1, NOTCH1, and CD133; the KI67/CD44 module comprises KI67 and CD44v1. Points are unconnected because donors are independent.

(C) Descriptive pathway-level summary of the broader historical Fluidigm panel at 10 µg/mL OPN. The complete donor-resolved numerical matrix was not preserved for every extended target; panel C is therefore descriptive and no gene-level inferential statistics are assigned to the extended panel.

### Supplementary Figure S4. WNT3A-conditioned-medium functional interaction analyses

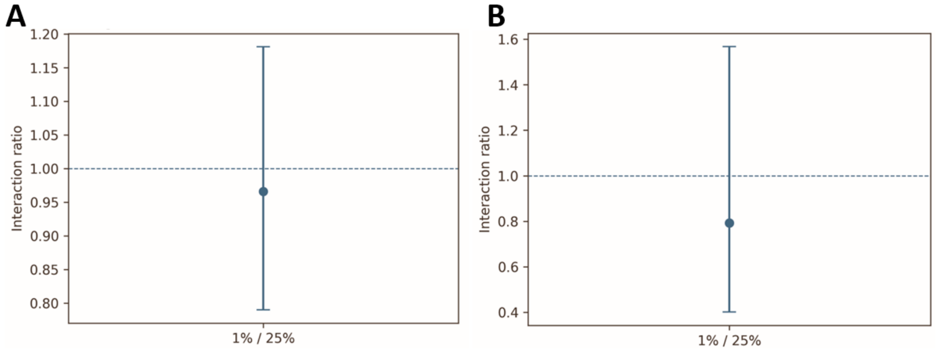


**Supplementary Figure S4.** Functional interaction analyses based on donor-level OPN/vehicle response ratios.

(A) For organoid area, the 1%/25% interaction ratio was 0.966 (95% CI, 0.790-1.181; P=.534).

(B) For EdU, the interaction ratio was 0.793 (95% CI, 0.401-1.567; P=.280). Neither analysis supports potentiation of OPN-dependent growth or proliferation at reduced WNT3A-conditioned-medium availability.

### Supplementary Tables

#### Supplementary Table S1. Clinical Metadata and Tissue Origin

| Culture | Age | Sex | Clinical/pathology context | Chemotherapy | Study role |
|---|---|---|---|---|---|
| NORG1 | 67 | M | Diverticulitis; mucosa without architectural disturbance or inflammatory infiltrate | No | Principal organoid donor |
| NORG2 | 74 | M | Caecal colonic adenocarcinoma; MMR-proficient/sporadic record | No | Principal organoid donor |
| NORG3 | 30 | F | Pelvic endometriosis involving the rectosigmoid junction | No | Principal organoid donor |
| NAF1 | Not available | Not available | Macroscopically noninflamed colonic tissue; metadata not available in supplied clinical table | Not available | Secretome NAF |
| NAF2 | 68 | M | Stage III well-differentiated colonic adenocarcinoma; noninflamed tissue used for NAF establishment | Yes | Secretome NAF |
| NAF3 | 47 | M | Stage III well-differentiated colonic adenocarcinoma; MMR-proficient | No/Not recorded | Secretome NAF |
| NAF4 | 53 | M | Rectal adenocarcinoma, stage III; rectosigmoid resection | No | Secretome NAF; conditioned-medium donor |
| NAF5 | 63 | F | Sigmoid adenocarcinoma, stage II | No/Not recorded | Secretome NAF |
| IAF1 | 33 | M | Crohn disease | No | Secretome IAF |
| IAF2 | 27 | F | Crohn disease | No | Secretome IAF; complementary fibroblast validation bank |
| IAF3 | 21 | M | Ulcerative colitis; severe acute colitis in supplied pathology wording | No | Secretome IAF |
| IAF4 | 48 | F | Crohn disease | No | Secretome IAF |

Note: NAF denotes fibroblasts established from macroscopically noninflamed tissue and should not be interpreted as uniformly healthy-donor fibroblasts. The validated secretome cohort comprises the 5 NAF and 4 IAF cultures listed above. Fibroblast SPP1 RT-qPCR validation was performed in a complementary 5-NAF/4-IAF cohort.

#### Supplementary Table S2. Culture-Medium Composition

| Component | Supplier/catalog | Condition | Final concentration / amount |
|---|---|---|---|
| Advanced DMEM/F12 | Invitrogen 12634-010 | DMEM/F12-HG base | 98% v/v |
| DMEM | Invitrogen 31966-047 | WNT3A-CM production | 89% v/v |
| MEM | Invitrogen 10370-047 | NAF4-CM production | 97.5% v/v |
| HEPES | Invitrogen 15630-056 | Relevant base media | 1% v/v |
| GlutaMAX | Invitrogen A1286001 | DMEM/F12-HG and NAF4-CM | 1% v/v |
| Fetal bovine serum | Invitrogen F2442 | WNT3A-CM / NAF4-CM | 10% / 0.5% v/v |
| L Wnt-3A cells | ATCC CRL-2647 | WNT3A-CM production | Producer cell line |
| NAF4 cells | Primary culture | NAF4-CM production | Producer cell culture |
| DMEM/F12-HG | Laboratory base | WENR / OPN-MCNAF | 48% / 24% v/v |
| WNT3A-conditioned medium | Laboratory produced | WENR / OPN-MCNAF | 48% / 24% v/v |
| WNT3A-conditioned medium | Laboratory produced | WNT perturbation experiments | 25% or 1% v/v |
| NAF4-conditioned medium | Laboratory produced | OPN-MCNAF | 48% v/v |
| N2 | Invitrogen 17502-048 | Organoid medium | 1% v/v, 1X |
| Nicotinamide | Sigma N0636 | Organoid medium | 10 mmol/L |
| B27 without vitamin A | Invitrogen 12587-010 | Organoid medium | 2% v/v, 1X |
| Human EGF | Invitrogen PHG0311 | Organoid medium | 8 nM |
| Human Noggin | PeproTech 120-10C-B | Organoid medium | 2 nM |
| Human R-Spondin 1 | R&D Systems 4645-RS-025 | Organoid medium | 40 nM |
| Gastrin | Sigma G9145-1MG | Organoid medium | 10 nM |
| SB202190 | Sigma S7067-5MG | Organoid medium | 10 μmol/L |
| PGE2 | Sigma P0409-1MG | Organoid medium | 10 nM |
| LY2157299 | Axon Medchem 1941 | Days 0-2 after passage | 500 nM |
| N-acetylcysteine | Sigma A9165-5G | Days 0-2 after passage | 1 mmol/L |
| Y-27632 | Sigma Y0503-1MG | Days 0-2 after passage | 10 μmol/L |
| Recombinant human OPN | PeproTech 120-35 | OPN experiments | 0.1-10 μg/mL |

### Supplementary Table S3. RT-qPCR Primers and Assay Validation

| Gene | Forward primer (5′→3′) | Reverse primer (5′→3′) | Efficiency / QC |
|---|---|---|---|
| KRT20 | CCCATCTCAGCATGAAAGAGTC | GGCGTTCCATGTTACTCCGA | 90%-110%; single melt-curve product validated |
| FABP1 | GCTGGGTCCAAAGTGATCCA | TATGTCGCCGTTGAGTTCGG | 90%-110%; single melt-curve product validated |
| CA2 | TGGACTGGCCGTTCTAGGTA | TCAGCACTCTTGCCCTTTGT | 90%-110%; single melt-curve product validated |
| MUC2 | ACTCCAACATCTCCGTGTCC | AGCCACACTTGTCTGCAGTG | 90%-110%; single melt-curve product validated |
| LGR5 | TGCTATTATTTCACCCCAATGCAT | GTTGGACGATAGGTCCAGCTTTA | 90%-110%; single melt-curve product validated |
| SOX9 | AGGAAGTCGGTGAAGAACGG | CTGGGATTGCCCCGAGTG | 90%-110%; single melt-curve product validated |
| HES1 | ACACGACACCGGATAAACCAA | AATGCCGCGAGCTATCTTTCT | 90%-110%; single melt-curve product validated |
| NOTCH1 | CCTCAACATCCCCTACAAG | ACACTTTGAAGCCCTCAG | 90%-110%; single melt-curve product validated |
| KI67 | TGCAGCAAGCACTTTGGAGA | CTTGACACACACATTGTCCTCAG | 90%-110%; single melt-curve product validated |
| CD44v1 | GCACAGACAGAATCCCTGCT | TCTTGCCTCTTGGTTGCTGT | 90%-110%; single melt-curve product validated |
| CD44v6 | TACCCCAGCAACCCTACTGA | CATCATTCCTATTGGTAGCAGGGA | 90%-110%; single melt-curve product validated |
| CD166 | TGGCAGTGGAAGCGTCATAA | CTCGTCTGCCTCATCGTGTT | 90%-110%; single melt-curve product validated |
| SNAI1 | CGAGTGGTTCTTCTGCGCTA | AGGGCTGCTGGAAGGTAAAC | 90%-110%; single melt-curve product validated |
| ITGA2 | GTGGCTTTCCTGAGAACCGA | GATCAAGCCGAGGCTCATGT | 90%-110%; single melt-curve product validated |
| ITGAV | GAGCGGGACCATCTCATCAC | GCAACTCCACAACCCAAAGT | 90%-110%; single melt-curve product validated |
| ITGB1 | CCTACTTCTGCACGATGTGATG | CCTTTGCTACGGTTGGTTACATT | 90%-110%; single melt-curve product validated |
| ITGB4 | GGTGCACAAGAAGAAGGACTG | AAAGCCCACCATGTGACCTC | 90%-110%; single melt-curve product validated |
| SPP1 | ACCTGACATCCAGTACCCTGA | ACGGCTGTCCCAATCAGAAG | 90%-110%; single melt-curve product validated |
| GAPDH | GAGAAGGCTGGGGCTCAT | TGCTGATGATCTTGAGGGCTG | 90%-110%; single melt-curve product validated |
| HPRT | CCTGGCGTCGTGATTAGTGA | GCAGCAAGACGTTCAGTCCT | 90%-110%; single melt-curve product validated |
| B2M | GTGCTCGCGCTACTCTCTC | TCTCTGCTGGATGACGTGAG | 90%-110%; single melt-curve product validated |

GAPDH, B2M, and HPRT stability across vehicle and OPN conditions was verified before combined normalization. Statistical testing was performed on DeltaCq-derived normalized quantities; fold change was used primarily for graphical display.

### Supplementary Table S4. Published Human Crohn Disease Data Used for External Contextualization

| Study / dataset | Cohort / compartment | Published observation | Relation to present data | Interpretive role |
|---|---|---|---|---|
| de Bruyn et al., 2018; GSE90607 | Matched primary ileal myofibroblasts from Crohn normal-appearing, inflamed, and stenotic regions | Region-associated fibroblast phenotypes persist ex vivo; stenotic cells show ECM/LOX/MMP abnormalities | Supports durable Crohn stromal remodeling; does not independently establish SPP1 direction | Fibroblast disease-state context |
| Mukherjee et al., 2023 | Full-thickness Crohn resections with single-cell profiling | Extensive fibroblast heterogeneity and intercellular signaling in Crohn strictures | Supports fibroblast heterogeneity and disease-associated stromal communication | Crohn stromal context |
| Kanke et al., 2022; GSE164985 | Treatment-naive Crohn colon; epithelial scRNA-seq | Reduced canonical LGR5+ representation with shift toward a crypt-top colonocyte state | Compatible with no LGR5 expansion but not a phenocopy of the OPN response | Disease-state dependence |
| Li et al., 2024; GSE266546 | Terminal ileum and ascending colon across active/inactive Crohn disease and controls | LCN2/NOS2/DUOX2-high epithelial cells expand in active Crohn disease | Supports inflammation-associated displacement of conventional epithelial states | Active Crohn epithelial context |
| Karakasheva et al., 2026 | Human Crohn tissue and patient-derived colonoids | Inflammatory secretory progenitor state with persistent epigenetic poising and cytokine-responsive expression | Conceptually concordant with persistent inflammation-responsive progenitor/immature states | Human colonoid context |
| Moraitis et al., 2025 | Injury models plus human macrophage/organoid coculture | Macrophage-derived NRG1 and SPP1 regulate injury-associated regenerative reprogramming | Supports OPN as one component of a multicue regenerative niche | Mechanistic context |

No de novo pseudobulk reanalysis of the public matrices is claimed. Published findings were used to contextualize the experimental results; publicly available datasets are cited in the manuscript reference list with accession numbers.

### Supplementary Table S5. Key Donor-Level Statistical and Sensitivity Summaries

| Analysis | Effect estimate | P / interval | Interpretation |
|---|---|---|---|
| Validated secretome OPN, 5 NAF vs 4 IAF | Cliff's delta=1.00; median ratio=3.64 | Exact Mann-Whitney P=.0159; BH q=.19 | Only analyte with complete positive IAF-versus-NAF separation; supported by SPP1 RT-qPCR |
| Crohn-only IAF1/2/3 | Complete separation from 5 NAF; median ratio=4.22 | Exact P=.0357; n=3 IAF | Direction and rank separation preserved in Crohn-derived subset |

| Analysis | Effect estimate | P / interval | Interpretation |
|---|---|---|---|
| Fibroblast SPP1 RT-qPCR | Approximately 10-fold higher mean expression in IAF than NAF | Two-sided unpaired Student t test, P<.05; 5 NAF vs 4 IAF | Orthogonal transcript-level validation |
| Day 9 organoid area | Geometric mean OPN/vehicle ratio approximately 1.10 | Exploratory P=.343 | No uniform donor-level growth effect |
| Day 9 EdU | Vehicle 12.3/10.9/7.5%; OPN 16.1/11.1/8.3% | Paired donor-level P=.282 | No uniform donor-level proliferative effect |
| WNT3A-CM x OPN, area | 1%/25% response ratio=0.966 | 95% CI 0.790-1.181; P=.534 | No supported interaction |
| WNT3A-CM x OPN, EdU | 1%/25% response ratio=0.793 | 95% CI 0.401-1.567; P=.280 | No supported interaction |
